# Three Muscle-Specific DAF-16/FOXO Transcriptional Targets Activated by Reduced Insulin/IGF-1 Signaling

**DOI:** 10.1101/2022.12.09.519372

**Authors:** Shifei Wu, Jinghua Gao, Yan Li, Charline Roy, Ying Wang, Ben Mulcahy, William Li, Sruthy Ravivarma, John Calarco, Wesley Hung, Mei Zhen

## Abstract

*C. elegans* insulin/insulin-like growth factor 1 signaling, IIS, affects diverse physiological processes through the DAF-16/FOXO transcription factor. Despite its ubiquitous presence in somatic cells, DAF-16’s effects exhibit prevalent tissue specificity as well as tissue crosstalk. This implies that tissue-specific DAF-16 transcriptional programs contribute to functional diversity of IIS. To further investigate this possibility, we sought muscle-cell-specific DAF-16 transcriptional targets. Using fluorescence-activated cell sorting to enrich for body wall muscle cells from young hermaphroditic adults, we compared the muscle cell mRNA transcriptomes under conditions of high and low IIS activity, with and without DAF-16. We further analyzed DAF-16a’s binding sites in muscle and intestine cells by chromatin-immunoprecipitation sequencing. Combined output of these analyses is 12 candidate DAF-16 targets enriched for muscle cells. Transcriptional and translational reporters for three out of the four top candidates - a secreted protein C54F6.5, a calcium-binding protein CEX-1/calexcitin, and a fatty acid metabolic enzyme MLCD-1/MCD - showed DAF-16-dependent activation specifically in body wall muscle cells. Notably, reporters for *C54F6.5* and *cex-1* exhibit DAF-16-independent, constitutive expression in non-muscle cells, explaining their low rank or absence from the DAF-16 target lists generated by whole-animal microarray or mRNA-sequencing analyses. These results highlight the need to examine FOXO targets in a cell-type-specific manner.

**Article Summary:** This study is relevant to those interested in functional specificity of signaling pathways. It describes a rigorous workflow to identify tissue-specific transcriptional changes activated by DAF-16/FOXO, a key effector of insulin signaling, its tissue-specific chromatin binding sites, and experimental validation of three previously unknown DAF-16 targets in body wall muscle cells. These findings highlight the intricacy of tissue-specific regulation exerted by a signaling pathway that is present and operates across tissues.

## Introduction

The insulin/insulin-like growth factor 1 (IGF-1) signaling (IIS) regulates a variety of physiological functions (Rask-Madsen and Kahn 2012), from developmental decision (Gottlieb and Ruvkun 1994; Morris et al. 1996; Kimura et al. 1997; Kaletsky and Murphy 2010), neural development (Man et al. 2000; Mielke et al. 2005; Plum et al. 2006; Chiu and Cline 2010; Klöckener et al. 2011; Nieto-Estévez et al. 2016), learning (Kodama et al. 2006; Tomioka et al. 2006; Lin et al. 2010; Oda et al. 2011; Chen et al. 2013; Tomioka et al. 2016), aging (Pan et al. 2011; Tank et al. 2011; Toth et al. 2012; Liu et al. 2013; Hahm et al. 2015; Li et al. 2019; Weng et al. 2024), longevity (Kenyon et al. 1993; Kimura et al. 1997; Ogg et al. 1997; Kaletsky and Murphy 2010), to stress response (McColl et al. 2010; Rasulova et al. 2021). Key components of the canonical IIS signaling pathway, consisted of the insulin and insulin-like ligands (INS), their respective transmembrane tyrosine kinase receptors, the cascade of phosphoinositide 3-kinase (PI3K) and serine/threonine kinase transducer of the receptor activity, and the effector of phosphorylation, the FOXO transcription factor, are conserved across animals (Kleemann and Murphy 2009; Piñero González et al. 2009; Kaletsky and Murphy 2010; Kenyon 2010). IIS and its physiological functions have been extensively investigated in *C. elegans*. Compared to other systems, *C. elegans* has significantly expanded genes encoding INS (Pierce et al. 2001: 20; Li et al. 2003: 28), while keeping a single, ancestral receptor DAF-2/InR for INS (Kenyon et al. 1993; Kimura et al. 1997). With remarkable complexity in their spatial and temporal expression pattern (Liu et al. 2014), 39 INS homologs (INS 1-37, INS-39 and DAF-28) either positively or negatively regulate the IIS activity (Chen et al. 2013; Hung et al. 2013; Murphy 2013; Fernandes de Abreu et al. 2014; Hung et al. 2014; Liu et al. 2014; Zhu and Chin-Sang 2024). An activated DAF-2 receptor activates a conserved PI3K (AGE-1) (Morris et al. 1996), PDK (PDK-1), and AKT (AKT-1 and AKT-2) (Paradis and Ruvkun 1998; Paradis et al. 1999) kinase cascade, leading to phosphorylation and cytoplasmic retention of the FOXO (DAF-16) transcription factor, inhibiting its transcriptional activity (Gottlieb and Ruvkun 1994; Ogg et al. 1997; Henderson and Johnson 2001; Lee et al. 2001; Libina et al. 2003; Lin et al. 2010) (Figure 1a). Consistent with IIS participating in diverse cellular functions, molecular components of the signaling cascade (from DAF-2/InR to DAF-16/FOXO) are expressed across cell-types (Aghayeva et al. 2021). Like the case for other signaling pathways, this raises a challenging mechanistic question: how does a single signaling cascade achieve its functional specificity?

**Figure 1.**
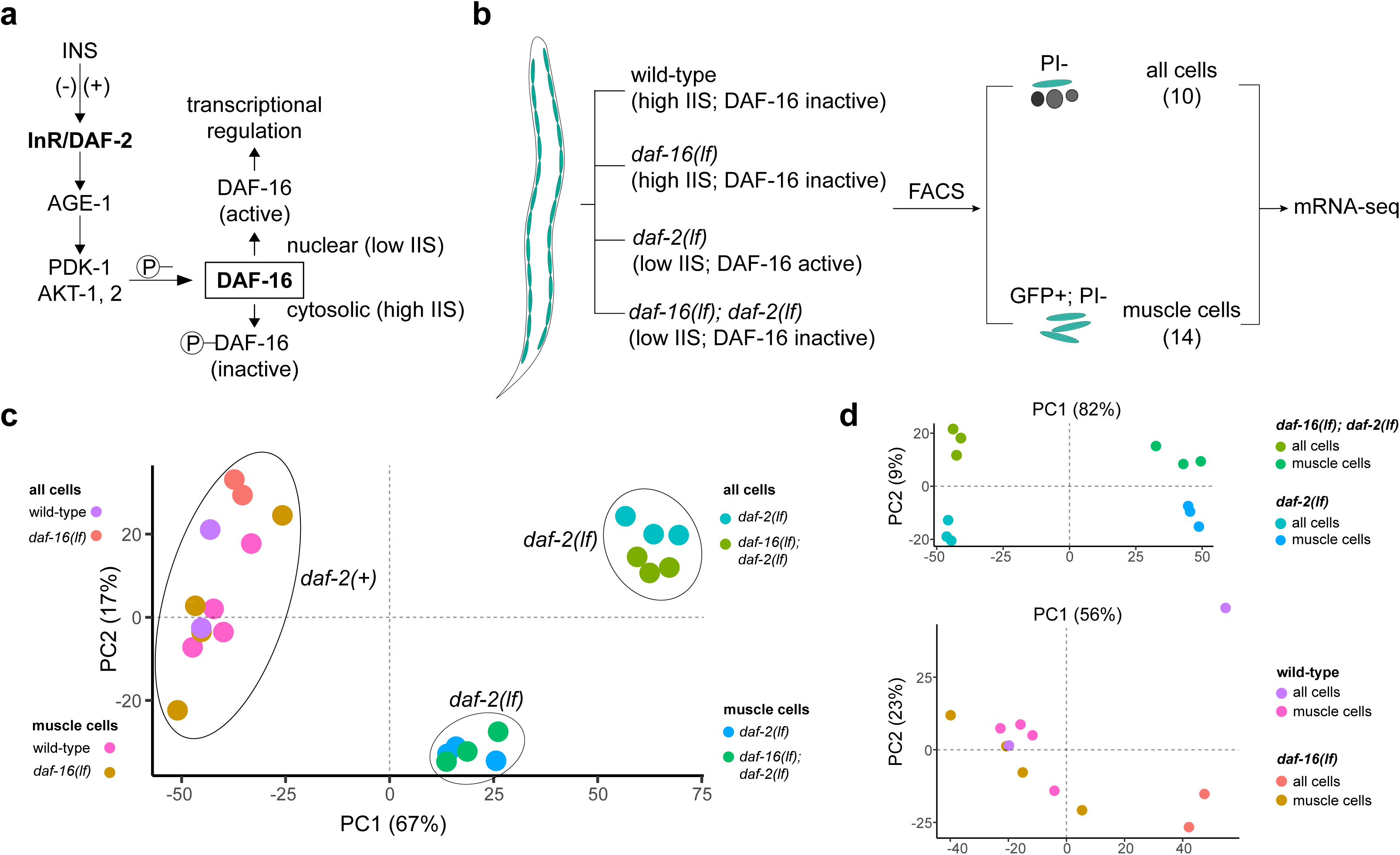
DAF-16 activity in different genetic backgrounds for mRNA-seq. (a) The effect of insulin/IGF-1 signaling (IIS) state on DAF-16 activity. A schematic of the IIS pathway in *C. elegans* is shown. Phosphorylation of DAF-16 sequesters the FOXO transcription factor from the nucleus, inhibiting DAF-16-directed transcriptional regulation. (b) Schematic of the FACS/mRNA-seq workflow for data collection. Animals of different genotypes (wild-type, *daf-16(lf)*, *daf-2(lf)*, *daf-16(lf); daf-2(lf)*) express GFP specifically in the body wall muscle cells. Age-matched young adults were collected, and cells were dissociated before being subjected to FACS. GFP+; PI- (propidium iodide negative) cells were sorted as ‘muscle cells’, whereas PI-cells were collected as ‘all cells’. Total mRNA was isolated for sequencing. (c) Principal component analysis (PCA) of the mRNA-seq datasets from 24 samples. (d) PCA of the mRNA-seq datasets from 12 *daf-2(lf)* samples (top) and 12 *daf-2(+)* samples (bottom).

Previous studies showed that functional specificity of *C. elegans* IIS results in part from the partial overlap of cells that produce different combination of INS (Murphy 2013). We and others further speculated that DAF-16 may define cell autonomous and non-autonomous functions of IIS by accessing different sets of transcriptional targets in a tissue-specific manner. Indeed, several processes regulated by IIS activity exhibit prominent tissue-specificity. A decreased IIS activity leads to increased adult lifespan (Kenyon et al. 1993; Kimura et al. 1997; Ogg et al. 1997; Kaletsky and Murphy 2010), and promotes an alternative ‘dauer’ developmental state that confers resistance to starvation and dehydration (Gottlieb and Ruvkun 1994; Morris et al. 1996; Kimura et al. 1997; Kaletsky and Murphy 2010). Both processes exhibit a strong requirement for an increased DAF-16 activity in the intestine. DAF-16’s presence in interneurons affect salt-associated learning (Kodama et al. 2006; Tomioka et al. 2006; Lin et al. 2010; Oda et al. 2011; Chen et al. 2013; Tomioka et al. 2016). In motor neurons, decreased IIS and subsequent activation of DAF-16 functions cell-autonomously to delay age-related decline in axon regeneration (Byrne et al. 2014). Even more intriguingly, IIS from both neurons and muscles contribute to age-dependent motility decline, but in opposite ways, at different phases of adulthood (Hahm et al. 2015; Roy et al. 2022).

DAF-16 regulates many critical processes in muscles, including synapse-targeting process (muscle arm) extension (Dixon et al. 2008), neuromuscular junction maturation (Hung et al. 2013), cell death prevention (Oh and Kim 2013), and heat-induced calcium overload resistance (Momma et al. 2017). Consistent with observations in other tissues, DAF-16 plays both autonomous and non-autonomous roles in muscle cells (Zhang et al. 2013). Cell-autonomous functions of DAF-16 might be explained by DAF-16’s transcriptional regulation of muscle-specific targets; on the other hand, inter-tissue communication might be mediated through its targets that either are secreted from muscles, or residing at the muscle surface to affect the development of other cells.

To date, efforts to identify DAF-16 targets have focused on measuring differential gene expression in the presence or absence of active DAF-16 in whole animals. Using either microarrays or mRNA-sequencing (mRNA-seq), these studies identified transcriptional changes relate to metabolism, oxidative-stress response, lifespan, and development (McElwee et al. 2003; Murphy et al. 2003; Lee et al. 2009; Kaplan et al. 2015; Hibshman et al. 2017). Towards the identification of tissue-specific DAF-16 targets, transcriptome profiling on all neuronal cells identified enriched transcripts; many were shown or speculated to participate in age-related memory decline or axon regeneration and learning (Kaletsky et al. 2016; Weng et al. 2024). However, comprehensive information on DAF-16 targets in a cell-type-specific manner is lacking.

Obtaining a list of high-confidence transcriptional targets of DAF-16 in a cell-type specific manner is a prerequisite for the mechanistic dissection on how IIS regulates diverse physiological processes in different cells. We want to contribute towards this goal by identifying DAF-16 targets specific for the body wall muscle cells. Our focus on muscle cells was motivated in part by the abundance and homogeneity of this tissue, and in part by the availability of raw DAF-16::GFP chromatin immunoprecipitation sequencing (ChIP-seq) datasets for the muscle cells (Kudron et al. 2018). We generated multiple FACS-sorted muscle cell bulk mRNA-seq datasets and combined them with available tissue-specific ChIP-seq (one muscle-specific and two intestine-specific) datasets to identify high-confidence muscle-enriched DAF-16 transcriptional targets. Reporters for top three candidates confirmed muscle-specific DAF-16-dependent expression. Overall, this study contributes towards understanding of how biology utilizes a limited number of signaling pathways to build richly diverse life forms.

## Materials and methods

### Strains and transgene generation

*C. elegans* strains were cultured on OP50-seeded NGM media (Brenner 1974) at 22.5°C (unless otherwise stated). *daf-2(e1370)* temperature-sensitive animals were cultured at the permissive temperature 15°C and shifted to the restrictive temperature for experiments (see below for details for each experiment). A list of strains and plasmids used in this study is provided in Table S1. Transgenic animals were generated by microinjection as described previously (Mello and Fire 1995). For plasmid-based reporters, stable extrachromosomal arrays of each reporter construct were generated and assessed for consistency of expression patterns. Afterwards, one stable extrachromosomal array of each reporter was crossed into *daf-16(lf)*, *daf-2(lf), daf-16(lf)*; *daf-2(lf)* for quantification purposes. Endogenous reporters for *gstk-2(syb5762)* and *cex-1(syb8040)* were generated by CRISPR-Cas9 (SunyBiotech) and outcrossed against N2 for at least four times.

### Transcriptional and translational reporter plasmid construction

*C54F6.5*: Two transcriptional (pJH4768, pJH5155) and two translational (pJH4815, pJH5133) reporters were examined. For transcriptional reporters, a promoter sequence (1225 bp upstream of its start codon and the first 106 bp of its coding sequence) was generated with PCR from wild-type genomic DNA. It was cloned into the Fire Vector that supplied a generic 3’ UTR derived from *unc-54* (a gift from Andrew Fire; Addgene kit # 1000000001) to create pJH4768. A *C54F6.5* 3’ UTR fragment (1.6 kb downstream of its stop codon) was generated with PCR from wild-type genomic DNA to replace the generic 3’ UTR of pJH4768 to generate pJH5155. For translational reporters, the remainder of C54F6.5 coding sequence was synthesized (IDT) and introduced into pJH4768 by Gibson Assembly to generate pJH4815, and into pJH5155 to generate pJH5133.

*cex-1*: *cex-1* 3’ UTR (1053 bp downstream of its stop codon) was used to replace the *unc-54* 3’ UTR in a previously described P*_cex-1_* mCherry reporter that contains 2862 bp upstream of its start codon (Ji et al. 2021) to generate the transcriptional reporter pJH4782. An endogenous translational reporter (*syb8040*) was generated by in-frame insertion of RFP::AID at the start codon.

*mlcd-1*: 949 bp upstream of its start codon and 990 bp downstream of its stop codon were included as its promoter and 3’ UTR sequence, respectively, to generate a transcriptional GFP reporter, pJH5134.

*gstk-2*: An endogenous transcription reporter *syb5762* was generated by inserting a *sl2-gfp* reporter after the C-terminal stop codon of the *gstk-2* genomic locus.

### FACS isolation of muscle cells and mRNA-seq preparation

Wild-type and *daf-16(lf)* worms with GFP-labeled muscles (Table S1, JAC127 and ZM9156) were synchronized and grown at 22.5°C throughout development until they became young adults. *daf-2(lf)* and *daf-16(lf); daf-2(lf)* animals with GFP-labeled muscles (Table S1, ZM10379 and ZM10380) were synchronized, grown at 15°C until L4 to prevent dauer formation during L1-L2 transition and molting arrest in larva stage afterwards exhibited by *daf-2(lf)*, and shifted to 22.5°C to activate DAF-16 for 8 hours. From these synchronized young adult worms, cells were extracted with SDS-DTT, dissociated with pronase, and filtered through a 20 μm cell strainer (Spencer et al. 2014). FACS analysis was performed to isolate GFP positive and propidium iodide (PI) negative muscle cells. A fraction of PI negative cells was ‘sorted’ without GFP gating as the “all cells” control sample. 20,000-40,000 sorting events were collected for each strain. We performed four independent replicates for the wild-type and *daf-16(lf)* ‘muscle cell’ datasets, and three independent replicates for the *daf-2(lf)* and *daf-16(lf); daf-2(lf)* ‘muscle cell’ datasets. RNA was extracted with TRIzol, cleaned up with DNAse I, and quality checked before being sent for sequencing using a protocols adapted from (Spencer et al. 2014). In parallel, aliquots of sorted muscle and all cells were plated, and muscle cell enrichment was confirmed.

### Analysis of mRNA-seq data

mRNA-seq raw reads were processed to produce transcript abundance and counts with Salmon 1.4.0. STAR and HTSeq were applied to validate estimated read maps from Salmon. Differential gene expression analysis was performed with DESeq2. GraphPad Prism and Pheatmap were used for visualizing abundance data.

### Analysis of ChIP-seq data

ChIP-seq raw data were obtained for XE1464 (ENCFF281NRH, ENCFF657MHD, ENCFF229JKH, ENCFF023XDP), ZM7247 (ENCFF004HMK, ENCFF018WCD, ENCFF889RBK), and ZM8745 (ENCFF288VDV, ENCFF846OUT, ENCFF963PAM) from a publicly available database (Kudron et al. 2018). To identify muscle and intestinal GFP::DAF-16a binding sites, these ChIP-seq raw reads were aligned with Bowtie 2 2.4.1 and further processed with SAMtools. Peaks were called with MACS2 for each replicate using the input ChIP-seq data without antibody enrichment for DAF-16 binding sequences. Peaks from two replicates were combined with IDR. ChIPseeker was used to identify genes that reside within 5 kb of the 5’ or 3’ of the peak, including the coding regions. ChromoMap was used to visualize binding sites.

### RT-qPCR

Animals were synchronized and cultured as for those used for mRNA-seq analyses. Total RNA was extracted from synchronized young adult worms using TRIzol and with the method described previously (Spencer et al. 2014). RT-qPCR was performed with Luna Universal One-Step RT-qPCR Kit from NEB on BioRad CFX96 Touch Real-Time PCR Detection System.

Each RNA sample was tested with gene-specific primers (see Table S2 for sequences). We used two housekeeping genes whose expression is not dependent on IIS as controls for expression. *pmp-3* has been previously described (Hoogewijs et al. 2008). *rpl-17* was chosen because its transcript level is high and consistent in all of our samples based on our mRNA-seq data. Each RT-qPCR experiment was performed using 4 different dilutions of extracted RNA from each sample. The experiment was repeated 3 times. The relative fold expression change was calculated using the 2^-ΔΔCT^ method. Specifically, at each concentration, the ΔCt between the gene of interest and *rpl-17* or *pmp-3* was calculated for all genes and for all samples. At each concentration, the ΔΔCt and thus fold expression change was then calculated between the wild-type sample and other samples. Statistical analysis for RT-qPCR was performed by two-way ANOVA with multiple comparisons between the mean of one condition with the mean of every other condition followed by Šidák correction.

### Confocal fluorescent microscopy

Stocks for imaging were kept at 15°C. A few adults were picked from each stock. Their first-generation progenies were cultured at 20°C until L4, from which transgenic worms were picked and shifted to 22.5°C (for Figure 4d, 5d, S2a, S4b) or 25°C (for Figure 4e, 4f, 5e, 6c) for one day until they became young adults. Culturing conditions for dauers and starved adults in Figure S2b, S2c, S3a, and S3b were as follows. For dauer experiments, worms raised at 15°C were grown at 25°C for at least 10 days to allow dauer formation. On the day of experiment, they were washed off the plate and treated with 1% SDS for 20 minutes to select for dauers. L2/L3 larvae born and raised at 15°C were used at controls in these experiments. For adult starvation experiments, ten young adult worms were hand-picked onto each plate and allowed to grow at 15°C. After the OP50 food was exhausted (∼7 days) worms were left on the plate for another 2-3 days. Young-adult-sized worms were picked for imaging. Well-fed young adults raised at 15°C were used as controls. For each set of comparison applying the same transgenic line, we applied the same culturing and imaging conditions and image processing procedures.

For imaging, live animals were placed with minimal M9 solution on dry 2% agarose pads covered with a coverslip. All images were acquired using a 20x or 60x objective on a Nikon Ti2 spinning disk confocal microscope equipped with an Andor Zyla VSC-07720 camera, with XY titled images taken with 15% overlap and Z-stacks were acquired at a step size of 2 μm (20x objective) or 1 μm (60x objective). Quantification was performed with ImageJ 2.3.0 (Schindelin et al. 2012) where we acquired the total and mean intensity for ROI (muscle cells, RIM) and the mean intensity of a background area in the same image. To correct for background for mean fluorescence, total ROI intensity was subtracted with the product of the background mean intensity and the size of muscle cell; Mean ROI intensity was subtracted with the background mean intensity. All images showed maximum projection. Statistical analysis for fluorescence intensity was performed by two-way ANOVA with multiple comparisons between the mean of one condition with the mean of every other condition followed by Šidák correction.

## Results

### 24 bulk mRNA-seq datasets to identify DAF-16-dependent transcriptional changes under high and low IIS states enriched in mature body wall muscle cells

To uncover muscle-enriched transcriptional changes, we started with a transgenic strain where only the body wall muscle cells are labelled by GFP, which we referred to as the wild-type background for mRNA-seq experiments henceforth. We used animals of four genetic backgrounds, all carrying the same transgene - wild-type, *daf-16 loss-of-function (lf)* mutants, *daf-2(lf)* mutants, and *daf-16(lf); daf-2(lf)* mutants (see Methods) - to model high (*daf-2(+)*) and low (*daf-2(lf)*) IIS activity, with and without DAF-16 (Figure 1b). We compared DAF-16-dependent transcriptional changes under the two IIS states separately.

Under the high IIS state, DAF-16 is maintained at the cytosol thus has low transcriptional regulation activity. To obtained DAF-16-dependent transcriptional changes, we compared the transcriptomes of *daf-2(+)* animals in the presence or absence of DAF-16, specifically, between wild-type animals and *daf-16(lf)* mutant animals raised as synchronized cultures under optimal laboratory culture conditions at 22.5°C and harvested 8 hours post the last larva molting.

Reduced IIS induces DAF-16’s nuclear translocation and elevation of its transcriptional activity. To obtain DAF-16-dependent transcriptional changes under the low IIS state, we compared the transcriptome between *daf-2(lf)* single mutant and *daf-16(lf); daf-2(lf)* double mutant animals. Because a full or partial functional loss of DAF-2 leads to arrest of or compromised reproductive development, we raised an allele of temperature-sensitive *daf-2(lf)* larvae at the permissive temperature (15°C) until they completed the last larva molt. They were further raised at the non-permissive temperature (22.5°C) for 8 hours to allow accumulation of DAF-16-dependent transcriptional changes (see Methods for further explanation for rationales and details).

To uncover muscle-enriched transcriptional changes, we dissociated these age-matched young adults to collect GFP-positive cells using fluorescence-activated cell sorting (FACS) (‘muscle cells’; Figure 1b). Three to four replicates, each with 20,000 to 40,000 sorting events, were obtained for each genotype. Control samples (‘all cells’; Figure 1b) were collected after each sample underwent dissociation and sorting but without GFP gating. These samples served as controls not only for muscle-enriched transcripts, but also, importantly, transcriptions that reflect cellular responses induced by the not-at-all subtle dissociation experimental manipulations. Overall, this left us 24 samples spanning four genotypes, all with both sorted ‘muscle cell’ and ‘all cell’ samples. We obtained on average 45 million reads per sample (ranged from 29 to 67 million reads) to ensure sufficient sequencing depth. They mapped to 20,800 protein-encoding genes, with at least one read in at least one dataset.

Our bulk assessment of the resulted sequencing datasets showed that they exhibit consistent differences, in an IIS state, DAF-16 activity, and cell-type dependent manner (Figure 1c, d). Principle component analyses (PCA) applied to all 24 datasets separated the 12 *daf-2(+)* samples from the 12 *daf-2(lf)* samples; the latter further segregated into two clusters by the cell-type (‘muscle cells’ versus ‘all cells’), all by the first two principal components (PC) that together account for 84% of the total variance (Figure 1c). Further PCA analyses, applied separately to the 12 *daf-2(+)* or 12 *daf-12(lf)* samples, showed that while 12 *daf-2(+)* samples (wild-type and *daf-16(lf)* samples) do not readily distinguish each other (Figure 1d, bottom panel), 12 *daf-2(lf)* samples segregated into 4 distinct clusters with two PCs that account for 91% of all variance – the 1^st^ PC predominantly separates them by being either muscle cells or all cells, and the 2^nd^ PC predominantly separates them by the presence or absence of DAF-16 (Figure 1d, top panel). Thus, these datasets contain sequencing information to distinguish between the high and low IIS states, and at the low IIS state, DAF-16-dependent transcriptional differences both unique to muscle cells and shared by all cells.

This evaluation warranted the subsequent application of DESeq2 analyses on these 24 datasets to identify specific sets of transcriptional changes that depend on DAF-16 activation. Henceforth, we refer all samples with a wild-type *daf-2* genotype (wild-type, *daf-16(lf)*) as at the high IIS state, and all samples with a *daf-2(lf)* mutation (*daf-2(lf)*, and *daf-16(lf); daf-2(lf))* as at the low IIS state. We refer samples both with no DAF-16 (*daf-16(lf)* and *daf-16(lf); daf-2(lf)*), and with low DAF-16 activity due to high IIS (wild-type *daf-2*) as samples with ‘inactive DAF-16’, whereas those with high DAF-16 activity, induced by low IIS (*daf-2(lf)*), as samples with ‘active DAF-16’.

### No prevalent DAF-16 transcriptional regulation in body wall muscle cells at high IIS state

First, we compared samples at high IIS state from wild-type animals and *daf-16(lf)* mutants. Consistent with a lack of distinct clustering, the DESeq2 analyses applied to ‘muscle cell’ samples did not reveal prevalent transcript changes in the presence or absence of DAF-16 (Figure 2a; Table 1). Briefly, among 20,800 genes that the sequencing reads mapped to, only 11 passed statistical tests for differentially expressed candidates (FDR < 0.05, n = 4). Among them, nine were of insufficient reads (less than 4 counts per million, CPM) in wild-type and *daf-16(lf)* muscle cells; only two, *ZK105.1* and *col-33*, have >10 CPM in one genotype and none 0 CPM in the other (Table 1). However, they are not among candidates at low IIS state (Figure 2b). We consider them unlikely to be DAF-16 targets. This negative outcome corroborates conclusions from previous mRNA-seq analyses applied to samples prepared from intact animals: there were no appreciable transcriptional differences between wild-type animals and *daf-16(lf)* mutants under normal culture conditions (Hibshman et al. 2017; Li et al. 2019).

**Figure 2.**
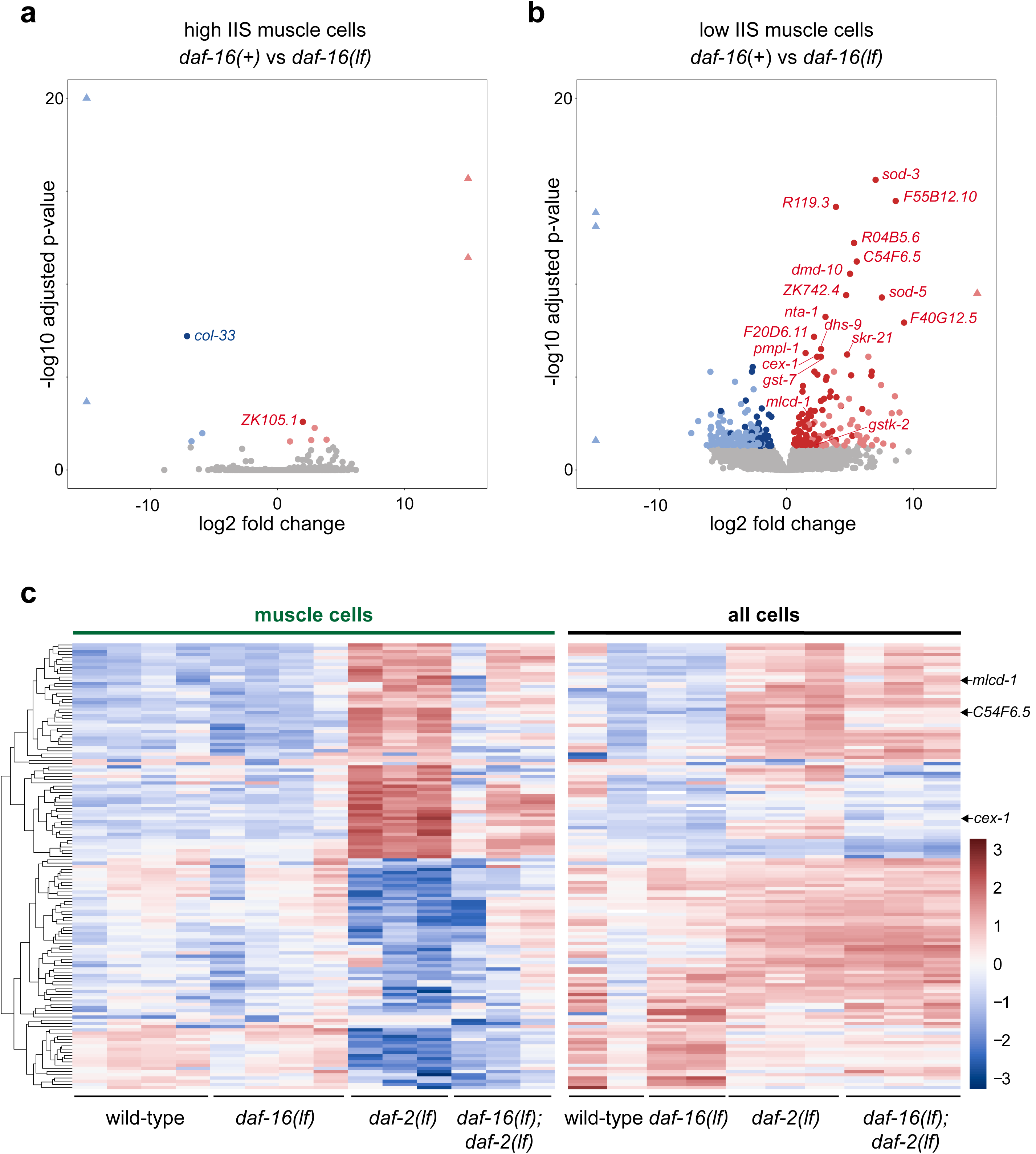
Differential gene expression analyses across transcriptomes of eight genotypes. (a, b) Volcano plots of differentially expressed genes (DEGs) for muscle cells at high IIS state, by comparing ‘muscle cell’ samples from wild-type and *daf-16(lf)* animals (a), for muscle cells at low IIS state, by comparing ‘muscle cell’ samples from *daf-2(lf)* and *daf-16(lf); daf-2(lf)* animals (b). Grey dots denote genes deemed unaffected by DAF-16 (FDR > 0.05). Red and blue dots denote genes upregulated and downregulated, respectively, by DAF-16 (FDR < 0.05). Red and blue dots with transparency denote those with <4 counts per million (CPM). Triangles denote values that exceed the scale at either axes. (c) Heatmap representation of transcript abundance of the 139 candidates for muscle-specific DAF-16-dependent transcriptional changes by mRNA-seq analyses. Transcript abundance in ‘muscle cells’ (left) and ‘all cells’ (right) were compared between four genetic backgrounds: wild-type, *daf-16(lf)*, *daf-2(lf)*, and *daf-16(lf); daf-2(lf)*. Transcript abundance was hierarchically clustered by similarities. Colors correspond to the z-score normalized over the average transcript abundance of each gene across all samples.

**Table 1.** DAF-16-dependent transcriptional regulation candidates at high IIS state.

|  | Salmon muscle CPM<br>(mean±SD, n=4) |  | STAR-HTSeq muscle<br>CPM (mean±SD, n=4) |  | DESeq2<br>based on Salmon |  |
| --- | --- | --- | --- | --- | --- | --- |
| Gene | wild-type | *daf-16(lf)* | wild-type | *daf-16(lf)* | log2<br>fold<br>change | adjusted<br>p-value |
| *col-33* | 0.16±0.15 | 30.53±46.51 | 0.17±0.16 | 34.47±52.29 | -7.10 | 6.32E-08 |
| *ZK105.1* | 15.86±7.03 | 4.06±2.11 | 18.27±6.84 | 4.59±1.98 | 2.00 | 2.58E-03 |
| *daf-16\** | 23.08±6.49 | 11.30±8.73 | 24.17±6.70 | 11.74±8.99 | 0.99 | 2.92E-02 |
| *Genes excluded due to low reads* |  |  |  |  |  |  |
| *gcy-3* | 0.02±0.03 | 0±0 | 0.02±0.02 | 0±0 | 17.27 | 2.11E-16 |
| *F23B2.10* | 0.01±0.02 | 0±0 | 0.03±0.04 | 0±0 | 15.99 | 3.82E-12 |
| *flr-4* | 0.86±1.15 | 0.05±0.03 | 0.89±1.24 | 0.08±0.07 | 3.87 | 2.34E-02 |
| *C55C3.3* | 3.57±2.17 | 0.49±0.40 | 3.93±2.51 | 0.51±0.39 | 2.93 | 5.47E-03 |
| *fpn-1.2* | 0.54±0.43 | 0.08±0.03 | 0.66±0.38 | 0.08±0.04 | 2.67 | 2.48E-02 |
| *C10F3.7* | 0.03±0.04 | 3.63±6.49 | 0.02±0.04 | 3.62±6.42 | -5.90 | 1.05E-02 |
| *srbc-15* | 0.01±0.02 | 2.17±2.99 | 0.01±0.02 | 1.86±2.41 | -6.75 | 2.86E-02 |
| *Y6B3B.1* | 0±0 | 0.07±0.12 | 0±0 | 0.06±0.11 | -17.86 | 1.67E-23 |
| *fbxb-33* | 0±0 | 1.75±3.48 | 0±0 | 0.01±0.01 | -28.14 | 2.15E-04 |

\*: Excluded from the candidate list of DAF-16-regulation events. Reduced reads resulted from the loss of transcription in the deleted genomic region in the *daf-16(mu86)* allele.

For our study, this outcome, which contrasts that at the low IIS state (below), born another critical significance. Because the dissociation and FACS-sorting process imposed acute cellular stress regardless of the genotype, this negative outcome showed that our experimental paradigm was able to maintain DAF-16 activity sufficiently low under normal growth conditions, and our analysis pipeline is sufficiently robust against noises induced by experimental manipulation.

In summary, we confirm that DAF-16’s transcriptional activity is low in muscle cells, where normal culture conditions maintain IIS at the high state.

### 195 candidates for DAF-16 regulation in body wall muscle cells at low IIS state

In contrast to samples at the high IIS state, and consistent with their distinct clustering, our DESeq2 analyses of the ‘muscle cells’ at the low IIS state revealed a substantial number of candidate DAF-16 targets. Transcript levels from 312 genes showed statistically significant differences between the *daf-2(lf)* and *daf-16(lf); daf-2(lf)* samples (FDR < 0.05, n = 3), 142 being upregulated and 170 being downregulated. This list warrants a further evaluation of DAF-16-dependent transcriptional regulation.

To identify the most consistent transcriptional changes induced by DAF-16 activation, we filtered this list against two more sets of transcriptomes with inactive DAF-16. Specifically, after we compared *daf-2(lf)* against wild-type, *daf-16(lf)* and *daf-16(lf); daf-2(lf)* ‘muscle cell’ samples, we kept those that showed statistical differences in all three comparisons. We reason that this step helps to reduce noise that may rise from stochastic compensation for a constitutive DAF-16 loss, and from the small number of replicates per genotype.

This leaves us 195 candidates - 88 upregulated and 107 downregulated - upon DAF-16 activation (Table 2; Table S3).

**Table 2.**
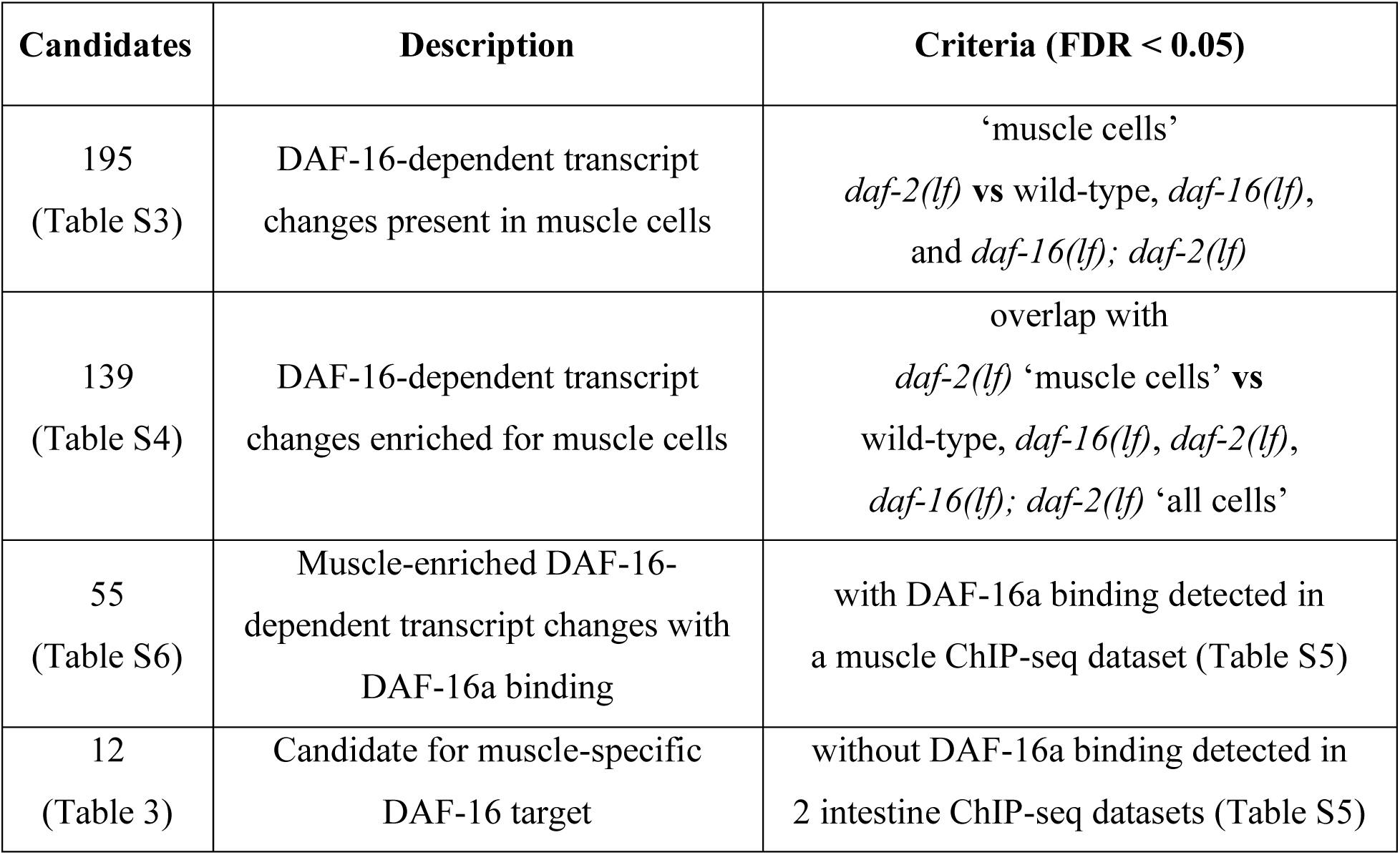
A summary of sequentially applied criteria for candidate DAF-16 targets.

| Candidates | Description | Criteria (FDR < 0.05) |
| --- | --- | --- |
| 195<br>(Table S3) | DAF-16-dependent transcript changes present in muscle cells | ‘muscle cells’<br>*daf-2(lf)* vs wild-type, *daf-16(lf)*,<br>and *daf-16(lf); daf-2(lf)* |
| 139<br>(Table S4) | DAF-16-dependent transcript changes enriched for muscle cells | overlap with<br>*daf-2(lf)* ‘muscle cells’ vs<br>wild-type, *daf-16(lf)*, *daf-2(lf)*,<br>*daf-16(lf); daf-2(lf)* ‘all cells’ |
| 55<br>(Table S6) | Muscle-enriched DAF-16-dependent transcript changes with DAF-16a binding | with DAF-16a binding detected in<br>a muscle ChIP-seq dataset (Table S5) |
| 12<br>(Table 3) | Candidate for muscle-specific DAF-16 target | without DAF-16a binding detected in<br>2 intestine ChIP-seq datasets (Table S5) |

### 139 candidates for body muscle cell-enriched DAF-16 regulation under a low IIS state

Among this list we sought candidates that are selectively regulated by DAF-16 in the body wall muscle cells. First, we compared the abundance of 195 candidates in active DAF-16 samples (*daf-2(lf))* that were sorted (‘muscle cells’) or not sorted (‘all cells’), and filtered 43 that exhibited similar abundances. Remaining 152 genes with either enrichment or depletion in muscle cells were further compared to two more sets of transcriptomes with inactive DAF-16 (wild-type and *daf-16(lf)*) from ‘all cells’. This resulted in a list of 139 genes (62 upregulated and 77 downregulated) to be preferentially affected by DAF-16 in body wall muscle cells (Figure 2c; Table 2; Table S4).

While a transcriptional regulation event shared by muscle and non-muscle cells does not exclude a muscle-specific function, we chose to focus on a smaller candidate list to further evaluate their candidacy as muscle-specific DAF-16 targets.

### 12 candidate DAF-16 muscle-specific targets under a low IIS state

Transcription factors affect a transcriptome directly, by binding the regulatory elements of a gene that leads to its activation or repression, as well as indirectly, such as by its primary targets modulating transcription in various ways. We sought DAF-16’s primary transcriptional targets by considering DAF-16-binding sites revealed by tissue-specific ChIP-seq analyses.

We previously generated a set of *daf-16(lf); daf-2(lf)* strains that each carry a single-copy transgene that expresses, from the same genomic locus, functional GFP::DAF-16a from the intestine cells or the muscle cells (Hung et al. 2014). They were used to generate the bulk of publicly available GFP::DAF-16a-ChIP-seq reads (Kudron et al. 2018). We previously chose the DAF-16a isoform because this, not the other (b, d/f) isoforms, was required in muscle cells for neuromuscular junction maturation (Hung et al. 2013). From these reads, we mapped 1,919 binding sites in the muscle-specific GFP::DAF-16a strain, and 955 and 2,422 binding sites in two independently derived intestine-specific GFP::DAF-16a strains (Table S5). ChIP-seq read peaks spanned widely across all six chromosomes (Figure 3a), consistent with the previous assessment that only a small fraction of DAF-16 binding events lead to transcriptional activation (Schuster et al. 2010).

**Figure 3.**
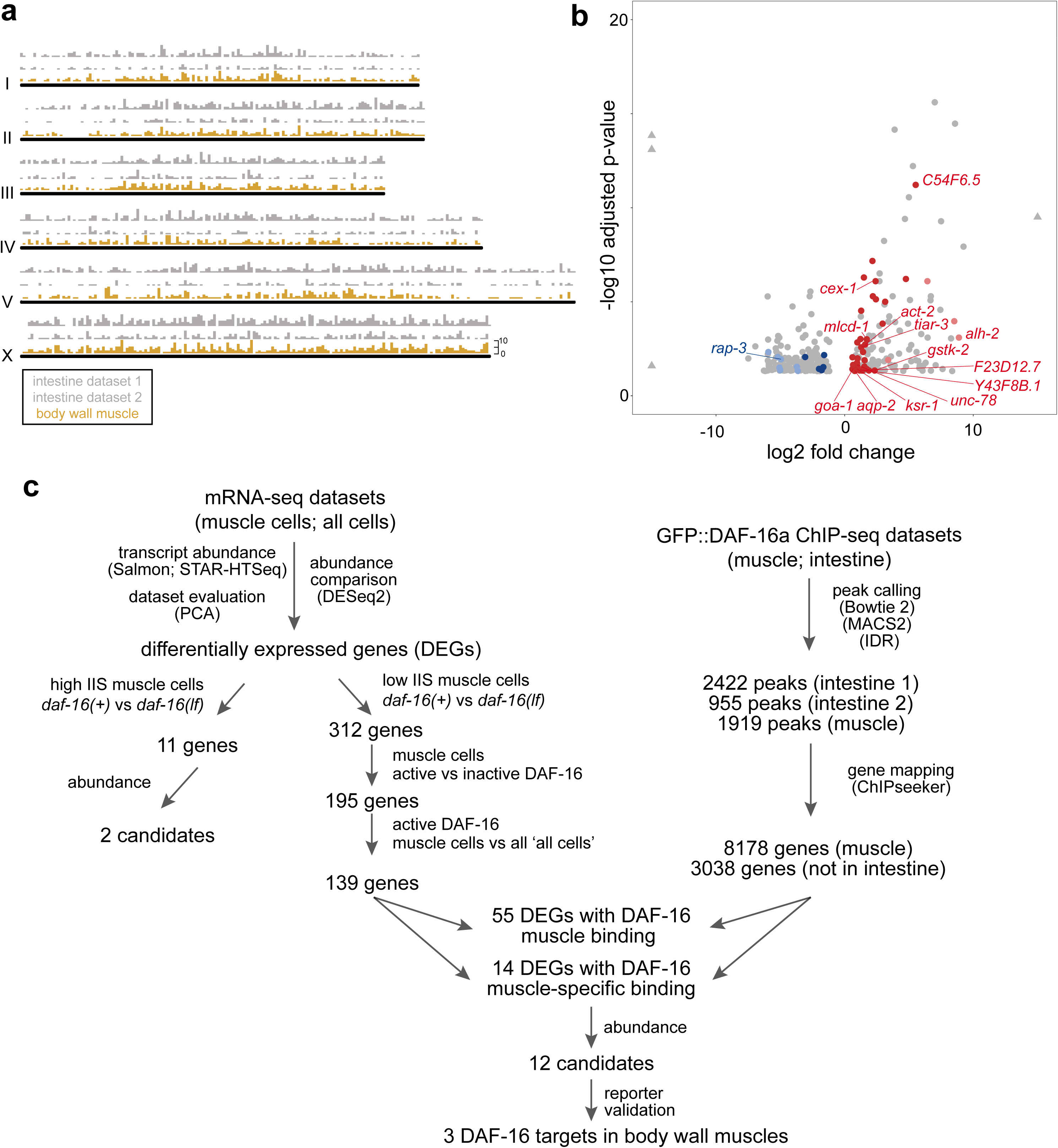
Combined DAF-16 ChIP-seq and comparative mRNA-seq analyses. (a) GFP::DAF-16a binding sites per chromosome, by two datasets for intestine ChIP-seq (grey) and one dataset for muscle ChIP-seq (brown). Peak numbers were binned for every 100,000 base pairs. (b) A volcano plot of 312 differentially expressed genes (DEGs) in muscle cells, by comparing ‘muscle cells’ samples from *daf-2(lf)* and *daf-16(lf); daf-2(lf)* animals. Grey dots denote genes without a muscle DAF-16 binding site within 5 kb flanking both sides of its coding region. Red and blue dots denote genes upregulated and downregulated, respectively, with a muscle DAF-16 binding site. Colored dots with denoted gene names have DAF-16 binding sites that are absent from both intestine ChIP-seq datasets. (c) A summary of our workflow to prioritize muscle-specific DAF-16 targets.

Among the 139 DAF-16 candidate targets, 55 showed at least one GFP::DAF-16a binding site within its genomic locus that begins at 5kb upstream of the first predicted or confirmed translational start site and ends at 5 kb downstream of the first predicted or confirmed stop codon. 43 are upregulated and 12 are downregulated by activated DAF-16 (Figure 3b; Table S6). Among them, 14 have the GFP::DAF-16a binding present in the muscle ChIP-seq dataset, but absent in both intestine ChIP-seq datasets. Removing those with low reads, we are left with 12 candidates, all being upregulated by activated DAF-16a (Figure 3b; Table 3).

**Table 3.**
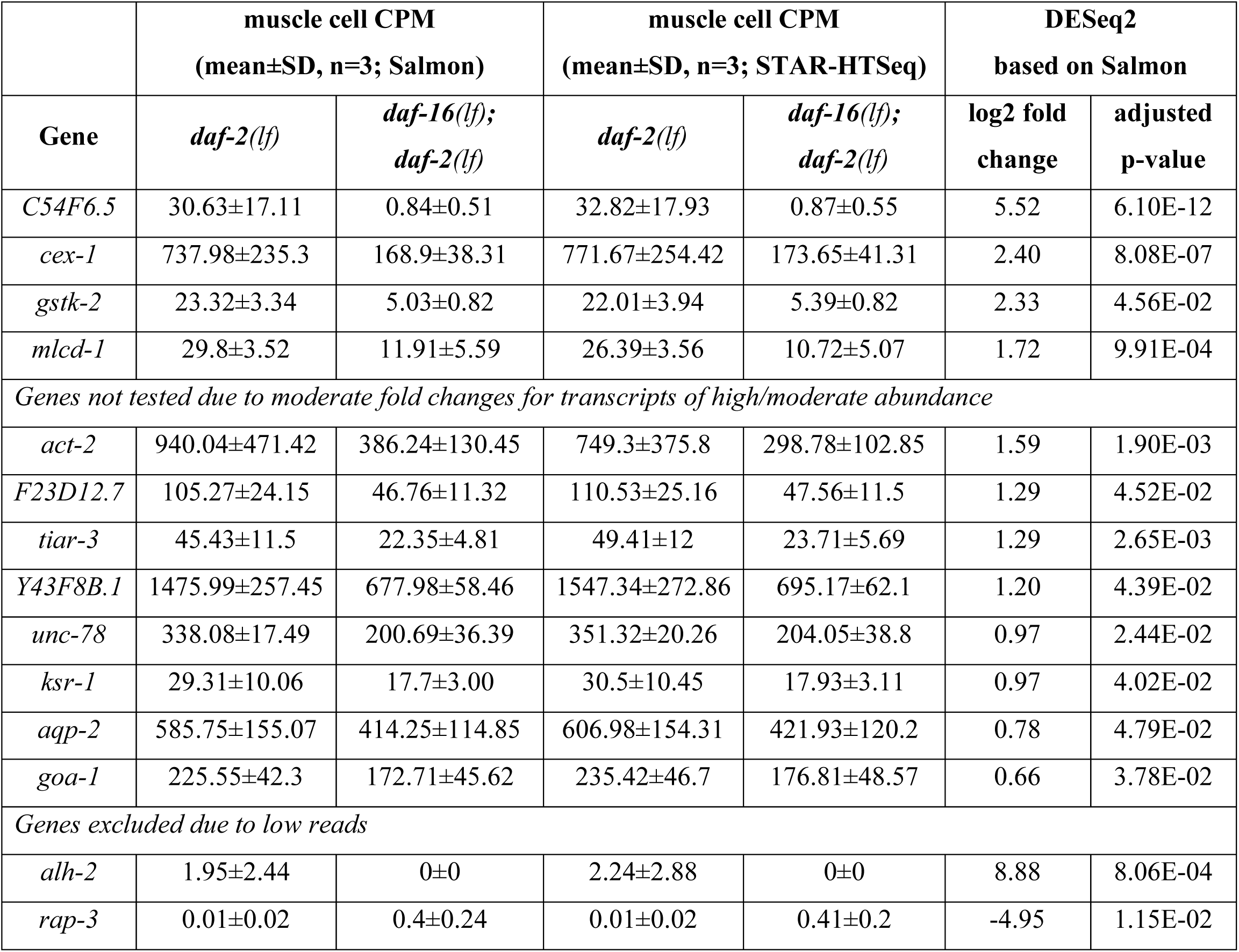
Twelve muscle-specific DAF-16 target candidates.

|  | muscle cell CPM<br>(mean±SD, n=3; Salmon) |  | muscle cell CPM<br>(mean±SD, n=3; STAR-HTSeq) |  | DESeq2<br>based on Salmon |  |
| --- | --- | --- | --- | --- | --- | --- |
| Gene | *daf-2(lf)* | *daf-16(lf);*<br>*daf-2(lf)* | *daf-2(lf)* | *daf-16(lf);*<br>*daf-2(lf)* | log2 fold<br>change | adjusted<br>p-value |
| *C54F6.5* | 30.63±17.11 | 0.84±0.51 | 32.82±17.93 | 0.87±0.55 | 5.52 | 6.10E-12 |
| *cex-1* | 737.98±235.3 | 168.9±38.31 | 771.67±254.42 | 173.65±41.31 | 2.40 | 8.08E-07 |
| *gstk-2* | 23.32±3.34 | 5.03±0.82 | 22.01±3.94 | 5.39±0.82 | 2.33 | 4.56E-02 |
| *mlcd-1* | 29.8±3.52 | 11.91±5.59 | 26.39±3.56 | 10.72±5.07 | 1.72 | 9.91E-04 |
| *Genes not tested due to moderate fold changes for transcripts of high/moderate abundance* |  |  |  |  |  |  |
| *act-2* | 940.04±471.42 | 386.24±130.45 | 749.3±375.8 | 298.78±102.85 | 1.59 | 1.90E-03 |
| *F23D12.7* | 105.27±24.15 | 46.76±11.32 | 110.53±25.16 | 47.56±11.5 | 1.29 | 4.52E-02 |
| *tiar-3* | 45.43±11.5 | 22.35±4.81 | 49.41±12 | 23.71±5.69 | 1.29 | 2.65E-03 |
| *Y43F8B.1* | 1475.99±257.45 | 677.98±58.46 | 1547.34±272.86 | 695.17±62.1 | 1.20 | 4.39E-02 |
| *unc-78* | 338.08±17.49 | 200.69±36.39 | 351.32±20.26 | 204.05±38.8 | 0.97 | 2.44E-02 |
| *ksr-1* | 29.31±10.06 | 17.7±3.00 | 30.5±10.45 | 17.93±3.11 | 0.97 | 4.02E-02 |
| *aqp-2* | 585.75±155.07 | 414.25±114.85 | 606.98±154.31 | 421.93±120.2 | 0.78 | 4.79E-02 |
| *goa-1* | 225.55±42.3 | 172.71±45.62 | 235.42±46.7 | 176.81±48.57 | 0.66 | 3.78E-02 |
| *Genes excluded due to low reads* |  |  |  |  |  |  |
| *alh-2* | 1.95±2.44 | 0±0 | 2.24±2.88 | 0±0 | 8.88 | 8.06E-04 |
| *rap-3* | 0.01±0.02 | 0.4±0.24 | 0.01±0.02 | 0.41±0.2 | -4.95 | 1.15E-02 |

Overall, this workflow applies progressively stringent criteria to narrow the candidacy of DAF-16 primary targets from an initially large list of transcriptional differences between the muscle cells harvested from *daf-2(lf)* and *daf-16(lf); daf-2(lf)* animals (Figure 3c: Table 2). Due to the limited number of replicates of mRNA-seq samples, insufficient replicates of ChIP-seq datasets as well as lack of raw reads of other tissue-specific ChIP-seq datasets, we took orthogonal approaches, described in Sections below, to further verify the candidacy of selected muscle-specific DAF-16 regulation.

### Validation of selected candidates with mRNA samples from non-transgenic strains

We ordered the 12 candidates, first by their fold changes between muscle samples with or without DAF-16 under a low IIS state, followed by their relative abundance (Table 3). Four top candidates are *C54F6.5*, *cex-1*, *gstk-2* and *mlcd-1*. Among them, all but *gstk-2* possess at least one DAF-16 binding element (DBE), a consensus DAF-16-binding cassette previously implicated in transcriptional regulation (Murphy et al. 2003; Schuster et al. 2010; Tepper et al. 2013), within or at the vicinity of identified DAF-16a binding sites (Figures 4c, 5c, 6b, S4b). All are ranked low for DAF-16-dependent transcriptional changes from whole-animal studies (Tepper et al. 2013).

**Figure 4.**
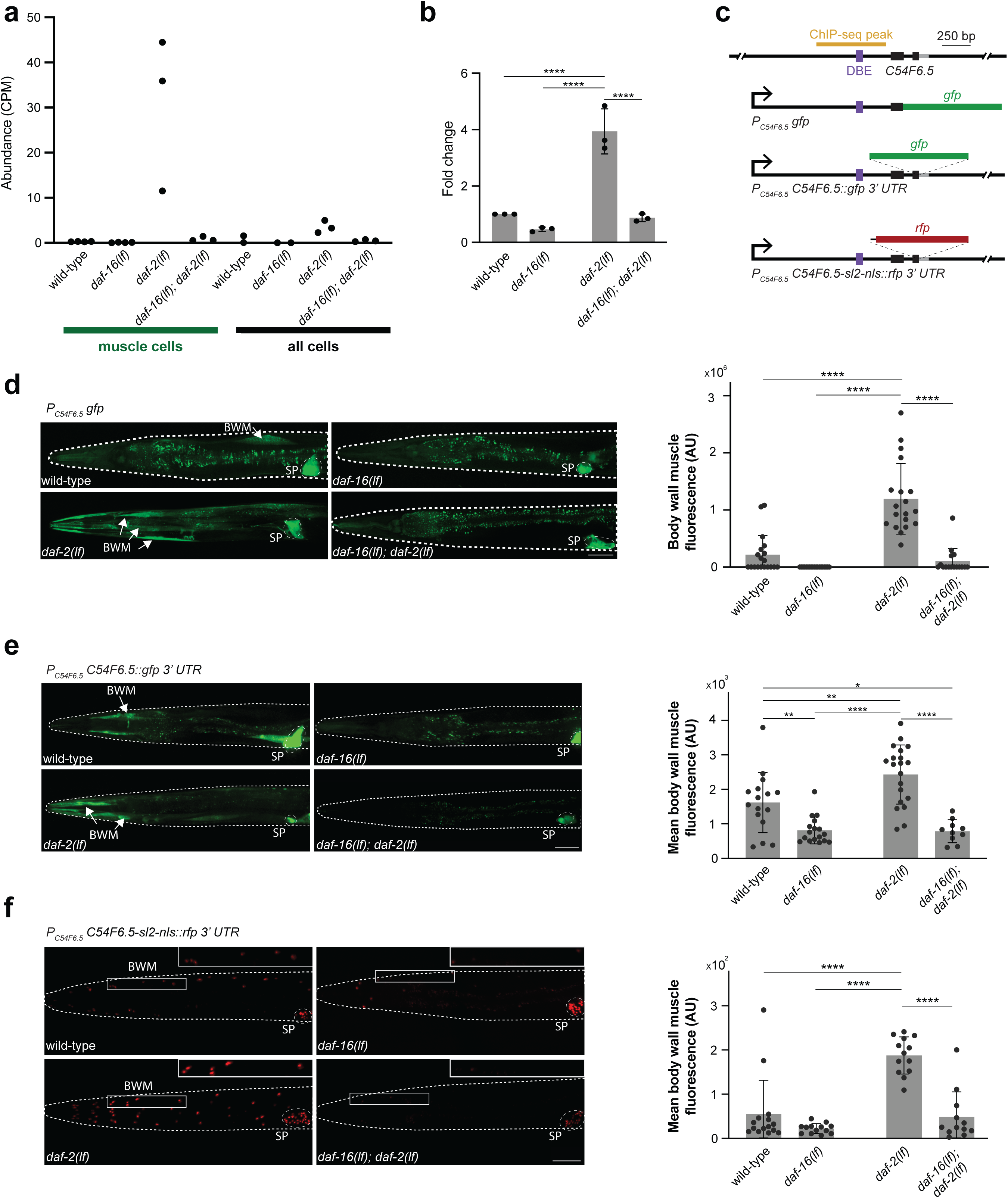
Muscle-specific upregulation of *C54F6.5* by DAF-16. (a) *C54F6.5* expression levels, measured as mRNA-seq read counts per million (CPM), across genetic backgrounds and cell types. (b) Fold changes of *C54F6.5* expression in respective genetic backgrounds, normalized against the wild-type level by RT-qPCR. Each data point represents the average of three cDNA dilutions (500pg, 5ng, and 50ng) from a single trial. n = 3 for all four genotypes. (c) The genomic structure of *C54F6.5* with its ChIP-seq peak, DAF-16 binding element (DBE), and reporter design labeled. (d) Representative confocal images and quantification of a *C54F6.5* transcriptional reporter in different genetic backgrounds. n = 19, 19, 19, 16. (e) Representative confocal images and quantification of a *C54F6.5* translational reporter with its endogenous 3’ UTR in different genetic backgrounds. n = 16, 17, 20, 10. (f) Representative confocal images and quantification of a *C54F6.5* transcriptional reporter with its endogenous 3’ UTR in different genetic backgrounds. n = 15, 13, 13, 12. BWM: body wall muscle. SP: spermatheca. All scale bars = 50 um. Non-significant comparisons were not labeled. **** p < 0.0001. ** p < 0.01. * p < 0.05. Error bars: standard deviation.

**Figure 5.**
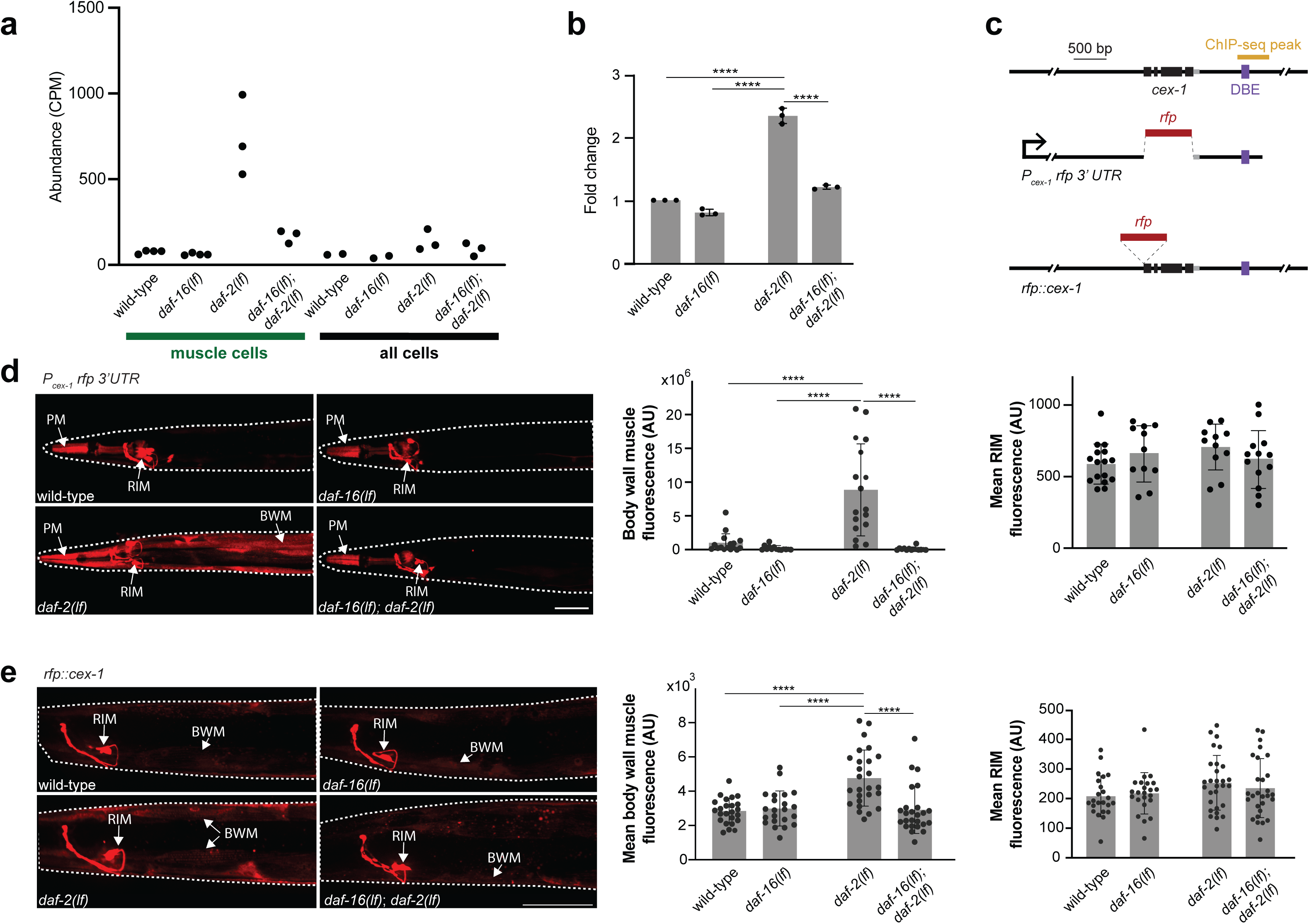
Muscle-specific upregulation of *cex-1* by DAF-16. (a) *cex-1* transcript abundance, measured as mRNA-seq read counts per million (CPM), across genetic backgrounds and cell types. Each dot represents the CPM from one sample. (b) Fold changes of *cex-1* expression in respective genetic backgrounds, normalized against the wild-type level by RT-qPCR. Each data point represents the average of three cDNA dilutions (500pg, 5ng, and 50ng) from a single trial. n = 3 for all four genotypes. (c) The genomic structure of *cex-1* with its ChIP-seq peak, DAF-16 binding element (DBE), and reporter design labeled. (d) Representative confocal images and quantification of a *cex-1* transcriptional reporter with its endogenous 3’ UTR in respective genetic backgrounds. n = 16, 14, 17, 14 animals for muscle. n = 16, 11, 11, 13 animals for RIM. (e) Representative confocal images and quantification of a *cex-1* endogenous reporter in respective genetic backgrounds. Each dot represents the image intensity for area of interest for one animal. n = 24, 22, 26, 25 animals for muscle. n = 23, 23, 29, 28 animals for RIM. BWM: body wall muscle; PM: pharyngeal muscle. All scale bars = 50 um. Non-significant comparisons were not labeled. **** p < 0.0001. Error bars: standard deviation.

**Figure 6.**
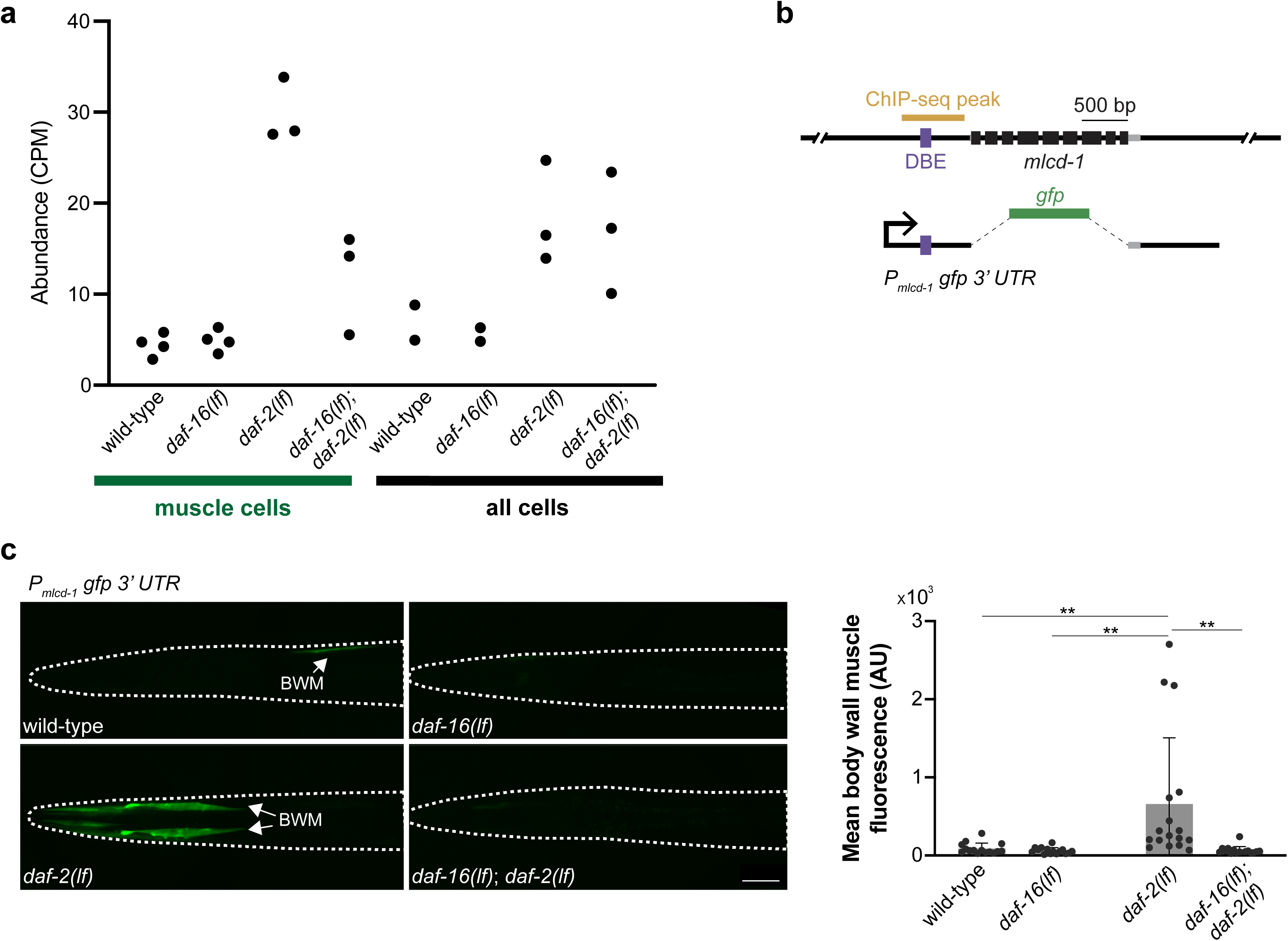
Muscle-specific upregulation of *mlcd-1* by DAF-16. (a) *mlcd-1* expression levels, measured as mRNA-seq read counts per million (CPM), across genetic backgrounds and cell types. (b) The genomic structure of *mlcd-1* with its ChIP-seq peak, DAF-16 binding element (DBE), and reporter design labeled. (c) Representative confocal images and quantification of an *mlcd-1* transcriptional reporter with its endogenous 3’ UTR in respective genetic backgrounds. n = 15, 15, 17, 16. BWM: body wall muscle. Scale bar = 50 um. Non-significant comparisons were not labeled. ** p < 0.01. Error bars: standard deviation.

Because our mRNA-seq datasets were obtained from dissociated cells, we wanted to first verify some candidates using RNA samples independently prepared from intact *C. elegans* samples and strains without transgenic markers. Using total mRNA prepared from synchronized and age-matched, non-transgenic wild-type (N2), *daf-2(lf)*, *daf-16(lf)*, and *daf-16(lf); daf-2(lf)* young adults, cultured in conditions similar to that of cell-sorting mRNA-seq experiments, we quantified the transcript level using RT-qPCR. We reason that upregulation of two high-abundance and high-fold change candidates, *C54F6.5* or *cex-1*, are the most likely be detectable in total RNA. Indeed, an upregulation of *C54F6.5* transcripts was also notable between our *daf-2(lf)* and *daf-16(lf); daf-2(lf) ‘*all cells’ samples (Figure 4a).

Our RT-qPCR analyses recapitulated DAF-16-dependent upregulation of *C54F6.5* and *cex-1* transcripts in *daf-2(lf)* mutants (Figure S1; Figure 4b; Figure 5b). Consistent with their rank, the transcript level of *C54F6.5* was the more robustly upregulated in active DAF-16 samples, similar to that of *sod-3*, a known pan-tissue DAF-16 target (Figure S1).

### DAF-16 upregulates multiple reporters for C54F6.5 in body wall muscle cells

*C54F6.5* is predicted (Blum et al. 2021) to encode a secreted 56-amino acid protein without functional characterization. With close to 0 read in all inactive DAF-16 muscle samples and high fold changes in active DAF-16 samples (Figure 4a; Figure 4b), we further examined its regulation with both transcriptional and translational reporters.

Immediately upstream of *C54F6.5* coding region resides a DAF-16a-ChIP-seq peak with DBE that is present only in the muscle datasets (Figure 4c, top panel). Several reporters with promoter sequences consisted of this region (Figure 4c, bottom panels; Methods) exhibited DAF-16-dependent, robust expression in body wall muscle cells or nuclei (Figure 4d-e, BWM) and DAF-16-independent expression in a non-muscle tissue, spermatheca (Figure 4d-e, SP). Consistent with *C54F6.5* being a secreted protein, translational reporters, where the full *C54F6.5* coding sequence fused with GFP, accumulated fluorescent signals in coelomocytes (Figure S2a), the scavenger cells that collect proteins in the body fluid (Fares and Greenwald 2001).

In body wall muscle cells, in the presence of DAF-16, both transcriptional reporters - one with endogenous 3’ UTR and another with a generic 3’ transcriptional enhancer - showed sporadic and weak signal in high IIS state (wild-type animals), but consistent and robust signals in low IIS state (*daf-2(lf)* mutant animals). Both sets of signals were eliminated by the absence of DAF-16 (*daf-16(lf)* and *daf-16(lf); daf-2(lf)*) (Figure 4d-f). Multiple lines of analyses thus consistently support the notion of a muscle cell-specific IIS regulation of *C54F6.5*.

### DAF-16 upregulates reporters for CEX-1/Calexcitin, a Ca^2+^ binding protein, in body wall muscle cells

CEX-1 is the *C. elegans* homologue of Calexcitin, a Ca^2+^ binding protein. *cex-1* exhibits low reads in inactive DAF-16 ‘muscle cells’, and high fold read increases in active DAF-16 samples (Figure 4a). Further verification of *cex-1* as a top candidate DAF-16 target by RT-qPCR (Figure 4b) was initially a surprise, because a *cex-1* promoter has long been used to drive constitutive gene expression in a single pair of interneuron (RIM) (Cohen et al. 2009; De Rosa et al. 2019). Our ChIP-seq analyses revealed a DAF-16a binding site with DBE at the 3’ UTR of *cex-1* that was only occupied in the muscle dataset (Figure 5c). Upon including *cex-1* 3’ UTR fragment with this site, a *cex-1* transcriptional reporter exhibited an expression pattern that corroborates the mRNA-seq and ChIP-seq results: a DAF-16-dependent upregulated expression in muscle cells (Figure 5d, BWM), and a DAF-16-independent constitutive expression in the RIM neuron (Figure 5d, RIM) and pharyngeal muscles (Figure 5d, PM). Consistent signals across four genotypes in RIM soma (Figure 5d, right panel) corroborated with a higher *cex-1* read counts in the ‘all cells’ samples by DAF-16-independent transcription activation.

Lastly, a *rfp::cex-1* translational reporter expressed from the endogenous genomic locus (Figure 5c) showed robust expression in the RIM neuron (Figure 5e, RIM), and low expression in body wall muscles comparable between wildtype, *daf-16(lf)*, and *daf-16(lf); daf-2(lf)* young adults (Figure 5e, BWM). DAF-16-dependent *rfp::cex-1* upregulation was observed in the body wall muscle cells of *daf-2(lf)* young adults (Figure 5e, *daf-2* panel, right panel). These results are consistent with a DAF-16-dependent *cex-1* transcriptional regulation specific for the body wall muscles and more prominent in low IIS state.

### DAF-16 upregulates a reporter for MLCD-1/MCD, a malonyl-CoA decarboxylase, in body wall muscle cells

*mlcd-1* encodes a fatty acid metabolic enzyme, malonyl-CoA decarboxylase (MCD). Ite exhibited overall lower transcript abundance than the other two candidates, but high fold changes (Figure 6a). A transcriptional reporter that consists of *mlcd-1* 5’ sequence with a DBE region and 3’ UTR (Figure 6b) showed body wall muscle-specific expression in *daf-2(lf)* young adults (Figure 6c, *daf-2* panel, BWM). The same transgene showed little signals in wild-type, *daf-16(lf)* and *daf-16(lf); daf-2(lf)* animals (Figure 6c). In our study, this reporter showed not only strong DAF-16-dependent expression, but also the highest specificity for body wall muscle cells.

### *C54F6.5*, *cex-1*, and *mlcd-1* reporter expression was altered in body wall muscle cells by physiological processes regulated by IIS

DAF-16 targets are likely to alter their expression in physiological states regulated by IIS. We qualitatively assessed how above reporters responded under two such physiological states, dauer formation and adult starvation.

For juvenile larvae, environmental stress such as food deprivation at high population density, triggers them to halt reproductive development and to enter dauer diapause (Fielenbach and Antebi 2008). This transition requires DAF-2-regulated transcriptional activation across tissues, and *daf-2(lf)* larvae activate dauer diapause even without food deprivation. We found that wild-type dauers (induced by poor environment), as well as *daf-2(lf)* dauers (induced by juvenile larvae at non-permissive temperatures), reporters for *C54F6.5* (Figure S2b and S2c, left panels), *cex-1* (Figure S3a, left panels), and *mlcd-1* (Figure S3b; left panels) all showed potent activation in body wall muscles BWM in Figure S2 and S3). The *cex-1* reporter was further activated in enteric muscles (AS, AD in Figure S3a, left panels).

Their responses to adult starvation, for which we surveyed both wildtype adults and *daf-2* adults raised at permissive temperature, showed a similar trend but less robust and consistent than in the dauer larvae. For example, while both *C54F6.5* transcriptional (Figure S2b, right panels) and translational reporters (Figure S2c, right panels) were activated in the body wall muscles of starved adults in comparison to fed adults, the transcriptional reporter exhibited a more prominent response and in starved *daf-2* adults. The *mlcd-1* transcriptional reporter similarly showed activation in both starved adults, but more prominently in *daf-2* adults (Figure S3b, right panels). The *cex-1* translational reporter showed the least response in starved adults (Figure S3b, right panels) when compared to dauer larvae (Figure S3b, left panels).

These qualitative assessment, especially the reporter’s activation in dauer larvae lends further support that these genes likely represent bona fide DAF-16 targets in body wall muscles.

### Ambiguity of remaining candidates

We consider the candidacy of all remaining genes to be low, because of their low reads, or a moderate fold change for transcripts with high or moderate reads (Table 3). We tested *gstk-2*, which ranked the highest among them (Table 3; Figure S4a), with an endogenous GFP reporter. Specifically, a cassette of splice leader 2 (*sl2*)-*gfp* inserted into the *gstk-2* locus drove co-transcription but separate translation of GSTK-2 and GFP (Figure S4b). It showed GFP expression in body wall muscles (BWM), pharynx cells (PH), and epithelial seam cells (SC) across wild-type, *daf-16(lf), daf-2(lf)*, and *daf-16(lf); daf-2(lf)* animals, without obvious differences in fluorescent intensity (Figure S4b). While we do not exclude experimental fallacies such as higher stability of the GFP reporter itself masking moderate differences at the protein level, we did not proceed further with this and remaining candidates.

## Discussion

A limited number of signaling pathways regulate numerous and diverse developmental processes in an organism- and cell-type-specific manner. IIS exerts physiological functions with tissue- and cell-specificity, and with both cell-autonomy and non-autonomy. Applying a rigorous workflow that reveals three new DAF-16 targets in body wall muscle cells, we contribute understanding towards the intricacy and mechanism of tissue-specific function exerted by a molecularly conserved and ubiquitously active signaling pathway.

### A small yield for tissue-specific DAF-16 targets

Combining mRNA-seq, ChIP-seq, RT-qPCR, and reporter analyses, we report three high-confidence DAF-16 targets primarily in the body wall muscle cells. Here we applied a series of increasingly stringent bioinformatic criteria that led to a small list of muscle-specific DAF-16 transcriptional targets (Figure 3c; Table 2). Reporter analyses for three of the four top candidates, all being overlooked in previous whole animal studies, lends support to this stringent approach.

Both negative and positive outcome from these analyses provided a few empirical guides to filter and enrich for differential gene expression events of physiological significance to follow through. Three targets constitute a minuscule fraction of the 139 genes that showed DAF-16-dependent, muscle-enriched transcript changes. We do not know how many primary DAF-16 targets drive the broader state of transcriptional changes, but we expected a lower representation of the primary transcriptional target, because a full transcriptome reflects the physiological state of cells.

However, this extremely low yield deserves further scrutiny, as described below.

### Limitation of this stringent approach

Our final candidate list (12 genes) is small, and three validated genes covered all high-confidence ones on the list. At this time, we do not know whether this short list reflects true physiology of muscle cells, where only a handful of molecular drivers underlie muscle-unique IIS regulation, or, high false negatives, where our stringent criteria led to excessive exclusion of DAF-16 transcriptional targets with functional relevance to muscle cells. Empirically, we favor the second possibility for several reasons.

First, by requiring both muscle-specific DAF-16 transcription regulation and muscle-enriched DAF-16a binding, we have biased against genes playing critical physiological functions at fast temporal dynamics or at low abundance.

Second, we have no criteria to objectively evaluate the coverage and consistency of the ChIP-seq reads. We noted that our sequential filtering during muscle-enriched transcriptomics analyses have maintained many high-confidence DAF-16 targets previously identified in whole-animal studies (Murphy et al. 2003; Tepper et al. 2013; Kaletsky et al. 2016) (Table S3, S4, S6), but many such candidates were removed when we applied the muscle ChIP-seq dataset as the first filter (Figure 3b, c). More specifically, imposing the presence of a binding site of muscle GFP::DAF-16a led to removal of ∼2/3 of candidates, which includes *sod-3*, a gold standard of the field and our positive control for RT-qPCR analyses (Figure S1). This raised concerns for missing targets due to the available muscle ChIP-seq reads.

Exclusion of isoforms other than DAF-16a in the ChIP-seq experiments may account for this concern; it is also possible that both muscle and intestine DAF-16 ChIP-seq experiments, being carried out as part of a systematic survey for transcription factors, were not as tightly controlled for developmental stages or genome coverage (Kudron et al. 2018), as for the focused mRNA-seq effort. Lastly, although we already broadened the region for gene mapping near the DAF-16a binding sites than regions considered for enrichment of the DBE sites in previous studies, regions to be explored remains an arbitrary criterion.

### Potential physiological functions of three DAF-16 targets

Physiological functions of three newly identified muscle DAF-16 targets are currently unknown, but we make a few speculations.

C54F6.5 is a small secreted protein. It has no apparent homolog outside the nematode specie by primary sequence alignment; however, we do not have reliable mean to predict structural similarity among small secreted proteins. Indeed, while amino acid sequences for many *C. elegans* neuropeptides revealed poor homologies, thus deemed to be ‘nematode-specific’, several have been shown to activate evolutionarily conserved GPCR signaling and to serve homologous physiological functions (Li 2008 Sep 25; Chen et al. 2022). C54F6.5 is one of many small secreted ‘nematode-specific’ proteins. These proteins are enriched for regulation upon exposure to pathogens and have been proposed to underlie the *C. elegans* immune response (Suh and Hutter 2012; Zhao et al. 2021). C54F6.5 might constitute a component of defense mechanisms contributed by muscle cells.

CEX-1 is homologous to calexcitin, a proposed Ca^2+^ sensor that regulates neuronal activity for learning (Nelson et al. 1996; Alkon et al. 1998; Nelson et al. 1999). With three Ca^2+^-binding EF-hand domains, it allows the cell to modulate its membrane excitability via the K^+^ channel and ryanodine receptor in a Ca^2+^-dependent manner (Alkon et al. 1998). Consistently, calexcitin was shown to bind ryanodine receptors in the sarcoplasmic reticulum (Gombos et al. 2001). In *C. elegans*, a bit ironically, there has been no functional studies on the *cex-1* gene itself, despite a fragment of its promoter being a popular tool to manipulate RIM, a neuron involved in associative learning, decision-making (Gordus et al. 2015; Jin et al. 2016; Sordillo and Bargmann 2021), and neuronal activation-induced stress response (De Rosa et al. 2019). CEX-1’s muscle up-regulation by DAF-16 promotes several hypotheses, for example, contribution to DAF-16-dependent effects on calcium homeostasis, motility, and health (Oh and Kim 2013; Roy et al. 2022; Momma et al. 2017).

MLCD-1, the *C. elegans* malonyl CoA decarboxylase (MCD) offers the clearest functional implication. MCD is abundantly expressed in the heart and skeletal muscles. Malonyl CoA is a potent inhibitor of carnitine palmitoyltransferase, an enzyme that transports long chain fatty acid into the mitochondria for ATP production. By breaking down malonyl CoA, MCD mobilizes fatty acid-driven energy production. Unlike the other two targets, the *mlcd-1* reporter is activated only in body wall muscles. Previous whole-animal RNA-seq datasets identified *mlcd-1* and other fatty acid metabolic enzyme-encoding genes to be upregulated by DAF-16 in context of longevity (Amrit et al. 2016), but our 139 muscle-enriched candidate DAF-16 targets do not include these other enzymes. MLCD-1 might modify the balance of energy storage and expenditure, instead of (or at least in addition to) simply boosting fatty acid oxidation in body wall muscle cells.

Muscle energy mobilization is particularly important for metabolism. We further speculate that DAF-16’s targeting of both CEX-1 and MCD-1 in muscle cells may reflect an orchestrated effort to increase muscle function as part of shared responses induced by reduced IIS signaling.

### Future directions

Future studies should address physiological functions of C54F6.5, CEX-1, and MLCD-1 in context of IIS signaling-related developmental and physiological processes. As mentioned in previous sections, behavioral and physiological consequences of single or combined knockout of these target genes, such as pathogen resistance, motility, and muscle physiology, under low and high IIS states serves as an excellent starting point. A requirement of either gene in respective phenotypes of *daf-2(lf)* mutants can then be further examined by standard molecular genetic manipulation, such as tissue-specific rescue or deletion.

Future studies should evaluate the validity of applying the ChIP-seq data to filter direct targets of transcription factors. This means that our 139 candidates from the mRNA-seq analyses should be independently evaluated for DAF-16-dependent transcriptional regulation. Ideally, a systematic, endogenous genomic tagging of these genes provides the needed assessment of required quality and coverage to incorporate information from future ChIP-seq datasets in search for cell-type specific and direct targets.

## Data Availability Statement

Strains underlined in Table S1, plasmids underlined in Table S1, and mRNA-sequencing data are available at CGC, Addgene, and GEO (GSE315486), respectively.

## Acknowledgments

We thank Kin Chan of the Network Biology Collaborative Centre Next-Generation Sequencing Facility (RRID: SCR_025385) at the Lunenfeld-Tanenbaum Research Institute for mRNA-seq. The facility is supported by the Canada Foundation for Innovation and the Ontario Government. We thank Arneet Saltzman and Tammy Lee for advice on mRNA-seq, ChIP-seq, and RT-qPCR. We acknowledge NIH support for CGC, where we deposited strains for distribution. We acknowledge WormBase (Sternberg et al. 2024) for support.

## Author contributions

M.Z. conceptualization; M.Z., W.H. supervision; S.W., J.G., W.H. experimental design, data collection and analyses; Y.L., C.R., Y.W., S.R., B.M., W.L., J.C. reagents and data collection; S.W., J.G. visualization; S.W., W.H., M.Z manuscript drafts; B.M. editing; M.Z. funding.

