## Supplementary figures and images for "Three Muscle-Specific DAF-16/FOXO Transcriptional Targets Activated by Reduced Insulin/IGF-1 Signaling"

### Figure S1

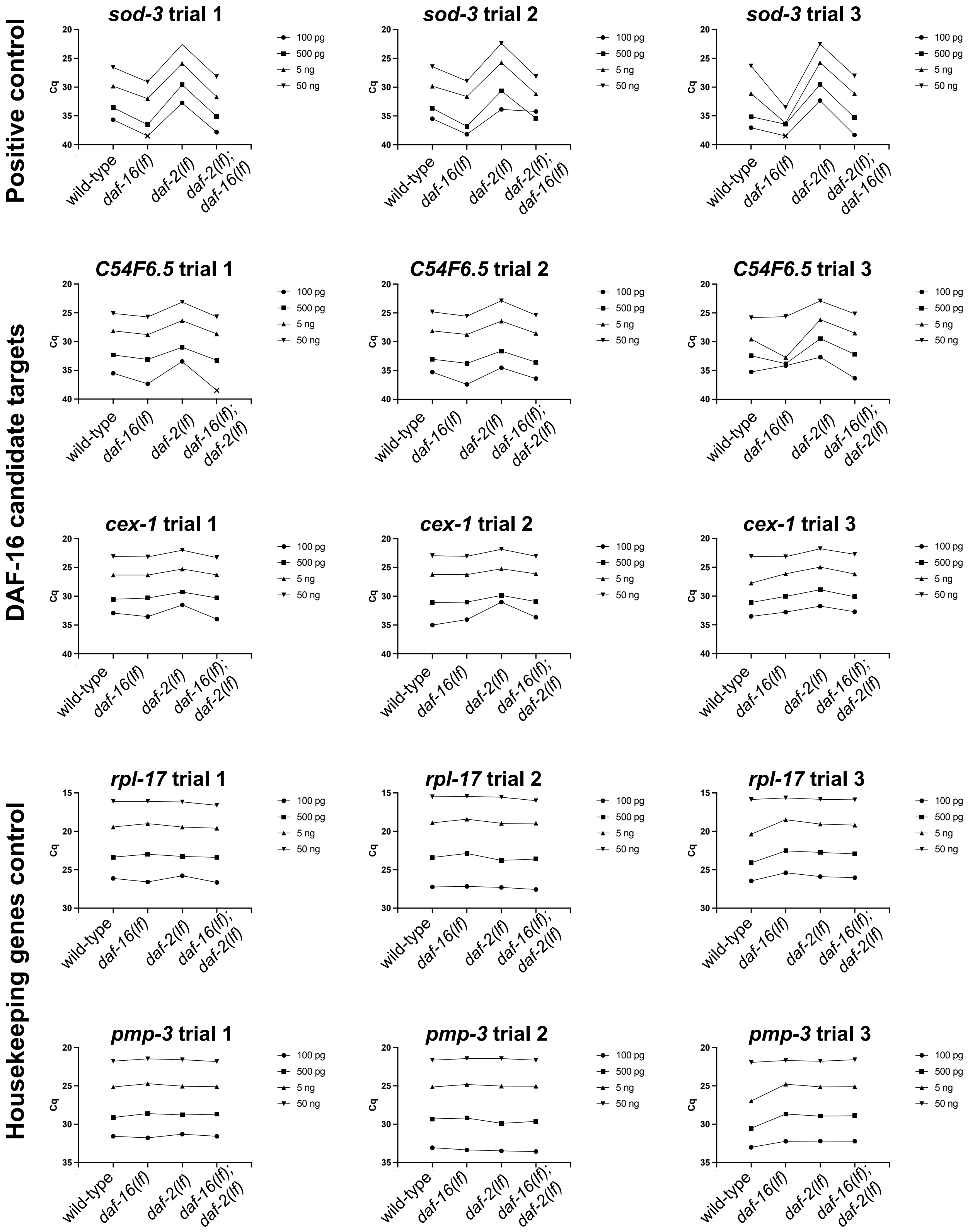

### Figure S2

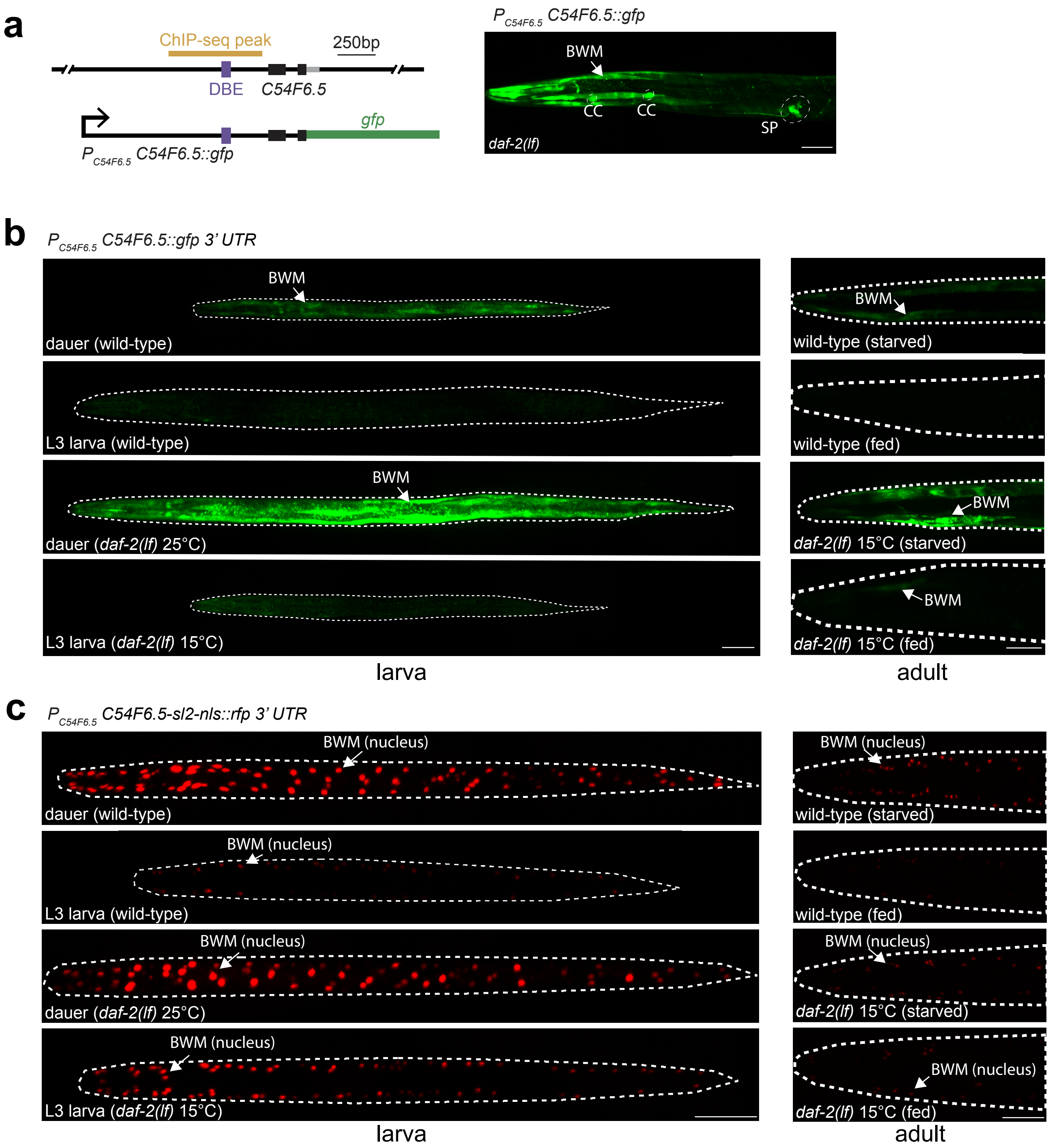

### Figure S3

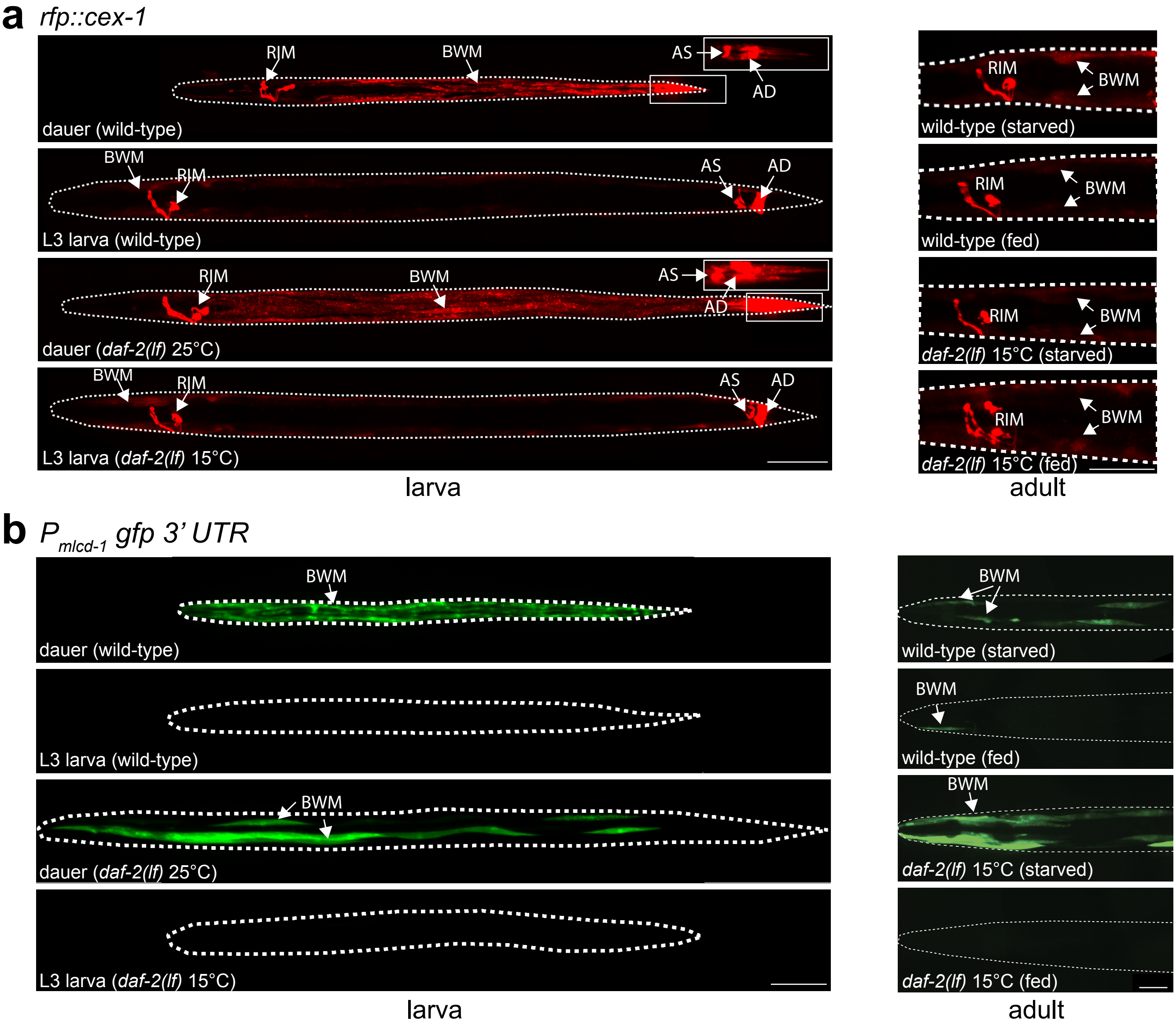

### Figure S4

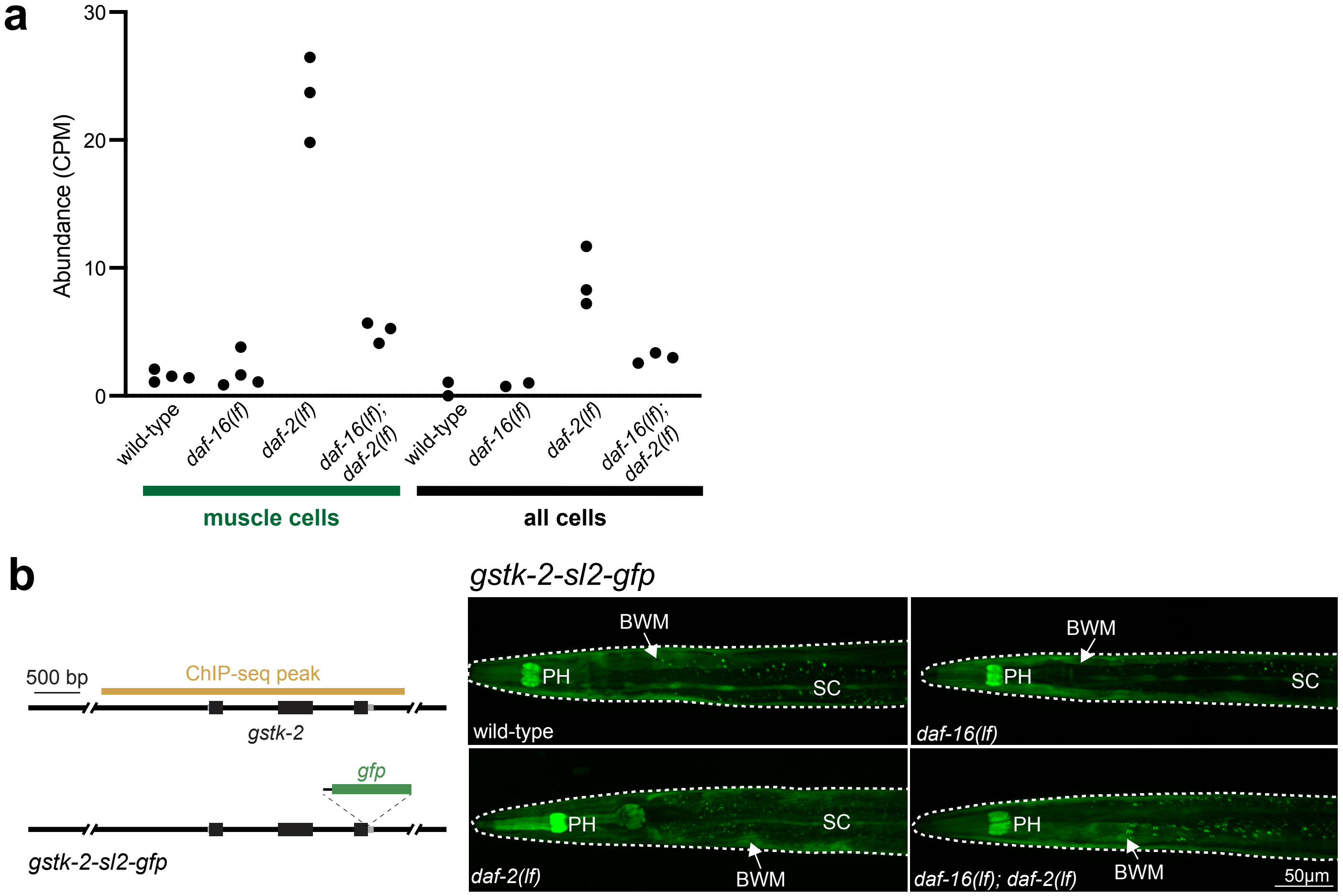
