## Supplementary material for "Three Muscle-Specific DAF-16/FOXO Transcriptional Targets Activated by Reduced Insulin/IGF-1 Signaling": Figure S1-S4 Legends

**Figure S1. Raw Cq values for 3 RT-qPCR replicates.**

RT-qPCR was performed for *C54F6.5*, *cex-1*, *sod-3* (positive control), *rpl-17* and *pmp-3* (housekeeping genes control) at different concentrations. *daf-2(lf)* showed consistent trends of transcript increase for *sod-3*, *C54F6.5*, and *cex-1*, but not for the *rpl-17* and *pmp-3* housekeeping genes.

**Figure S2. Reporters of *C54F6.5*, a secreted protein, are upregulated in body wall muscle cells in dauers and starved adults.**

(a) Genomic structure of *C54F6.5* with its ChIP-seq peak, DAF-16 binding element (DBE), and a translational reporter design labeled. A representative confocal image of the *C54F6.5* reporter showed signals in body wall muscle cells (BWM), coelomocytes (CC), and spermathecae (SP). (b) Representative confocal images of a *C54F6.5* translational reporter in dauers and respective non-dauer control larvae (left panels) and starved and respective fed control adults (right). BWM: body wall muscle. (c) Representative confocal images of a *C54F6.5* transcriptional reporter in dauers and respective non-dauer control larvae (left panels) and starved and respective fed control adults (right). BWM: body wall muscle. All scale bars = 50 um.

**Figure S3. Upregulation of *cex-1* and *mlcd-1* reporters in body wall muscle cells in dauers and starved adults.**

(a) Representative confocal images of a *cex-1* endogenous reporter in dauers and respective non-dauer control larvae (left panels), and starved and respective fed control adults (right panels). BWM: body wall muscle; AS: anal sphincter muscle; AD: anal depressor muscle. (b) Representative confocal images of an *mlcd-1* transcriptional reporter in dauers and respective non-dauer control larvae (left panels), and starved and respective fed control adults (right panels). All scale bars = 50 um.

**Figure S4. An endogenous *gstk-2* translational reporter does not show DAF-16-dependent expression level changes.**

(a) *gstk-2* expression levels, measured as mRNA-seq read counts per million (CPM), across genetic backgrounds and cell types. (b) Genomic structure of *gstk-2* with its ChIP-seq peak without a DBE binding site and the *gstk-2-sl2-gfp* reporter design labeled (left panel).

Representative confocal images of an endogenously expressed *gstk-2-sl2-gfp* reporter showed a similar expression in the body wall muscle (BWM), pharynx (PH), and seam cells (SC) across wild-type, *daf-16(lf)*, *daf-2(lf)*, and *daf-16(lf); daf-2(lf)* animals. Scale bar = 50 um.
